# Genetic and pharmacological correction of impaired mitophagy in retinal ganglion cells rescues glaucomatous neurodegeneration

**DOI:** 10.1101/2025.02.13.638142

**Authors:** Prabhavathi Maddineni, Bindu Kodati, Balasankara Reddy Kaipa, Karthikeyan Kesavan, J Cameron Millar, Sam Yacoub, Ramesh B Kasetti, Abbot F Clark, Gulab S Zode

## Abstract

Progressive loss of retinal ganglion cells (RGCs) and degeneration of optic nerve axons are the pathological hallmarks of glaucoma. Ocular hypertension (OHT) and mitochondrial dysfunction are linked to neurodegeneration and vision loss in glaucoma. However, the exact mechanism of mitochondrial dysfunction leading to glaucomatous neurodegeneration is poorly understood. Using multiple mouse models of OHT and human eyes from normal and glaucoma donors, we show that OHT induces impaired mitophagy in RGCs, resulting in the accumulation of dysfunctional mitochondria and contributing to glaucomatous neurodegeneration. Using mitophagy reporter mice, we show that impaired mitophagy precedes glaucomatous neurodegeneration. Notably, the pharmacological rescue of impaired mitophagy via Torin-2 or genetic upregulation of RGC-specific Parkin expression restores the structural and functional integrity of RGCs and their axons in mouse models of glaucoma and *ex-vivo* human retinal-explant cultures. Our study indicates that impaired mitophagy contributes to mitochondrial dysfunction and oxidative stress, leading to glaucomatous neurodegeneration. Enhancing mitophagy in RGCs represents a promising therapeutic strategy to prevent glaucomatous neurodegeneration.

**Graphical Abstract.**
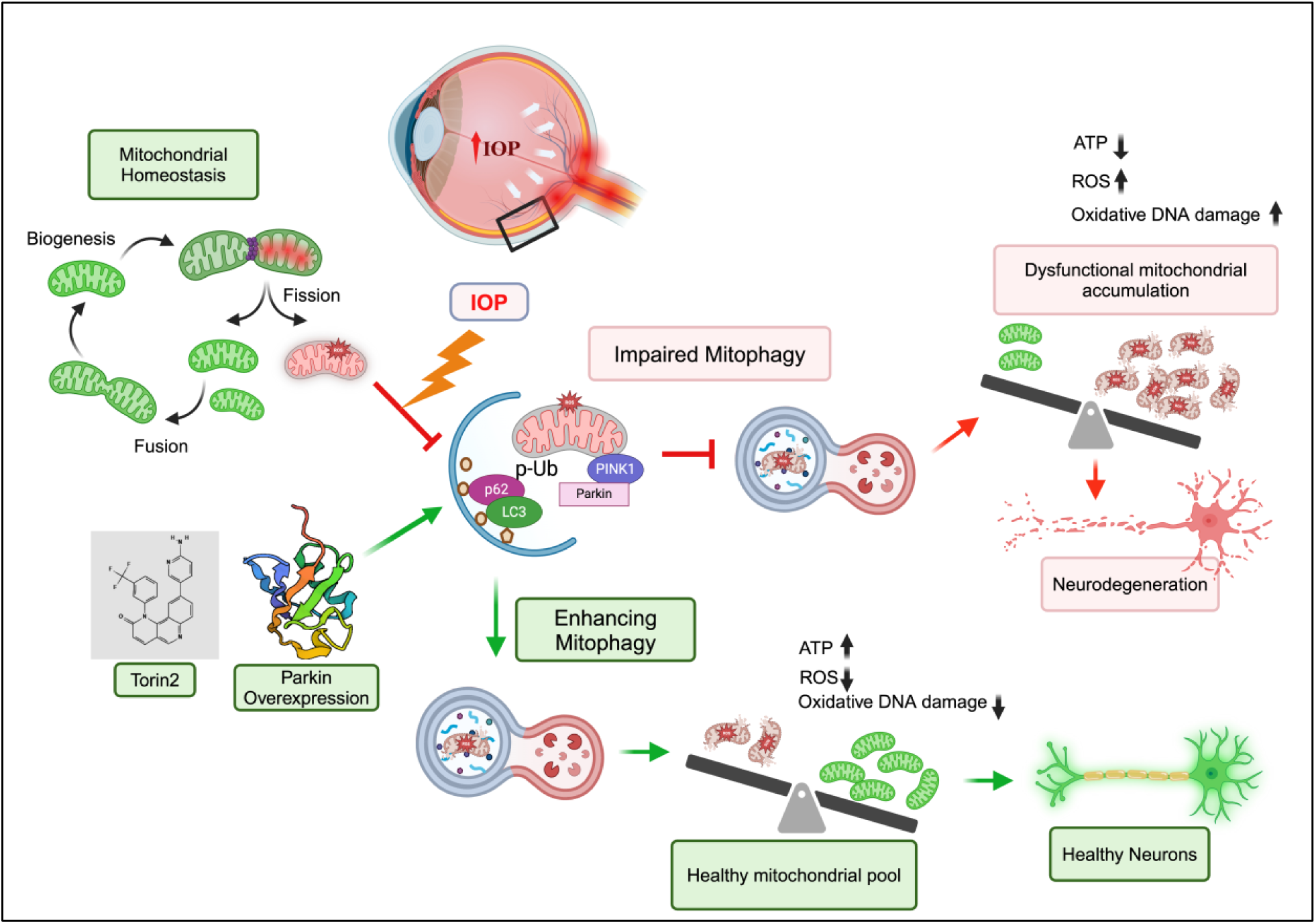

## Introduction

Glaucoma is a multifactorial neurodegenerative disease characterized by progressive loss of retinal ganglion cells (RGCs) and degeneration of the optic nerve (ON) axons, leading to irreversible vision loss. Glaucoma affects nearly 80 million people worldwide and is estimated to reach 111.8 million by 2040, making it the second leading cause of blindness in the world (1–3). Despite its high prevalence, the underlying pathological mechanisms responsible for glaucomatous neurodegeneration remain unclear. Age and elevated intraocular pressure (IOP) due to trabecular meshwork (TM) dysfunction are the major risk factors for the onset and progression of glaucoma (4, 5). Chronic elevation of IOP induces pathological changes at the optic nerve head (ONH), leading to RGC degeneration. These changes are accompanied by impaired axonal transport, glial/astrocyte activation, increased metabolic stress, and neuroinflammation. The primary therapeutic approach to the management of glaucoma is focused on reducing IOP. However, vision loss continues to progress in some glaucoma patients despite adequate therapeutic control of IOP, suggesting that there are other factors that contribute to the progression of the disease (6–8). Furthermore, long-term follow-up clinical studies have also shown that current IOP-reducing therapeutic approaches are insufficient, with approximately 40% to 60% of patients experiencing glaucoma progression after seven to eight years (9–11). Therefore, identifying and targeting IOP-induced early events of glaucomatous neurodegeneration is crucial for developing effective glaucoma treatments.

Degeneration of RGC soma and axons is the pathological hallmark of glaucoma. RGCs form the innermost layer of the neural retina and are the only retinal neurons that communicate directly with the brain. RGCs receive vast visual information from the photoreceptors via retinal interneurons (including horizontal, bipolar, and amacrine cells) and relay visual information in the form of action potentials to the visual centers of the brain. Therefore, RGCs rely mainly on mitochondria to support their high bioenergetic requirements to maintain their complex structure and functional integrity (12). In RGCs, mitochondria redistribute continuously across dendrites, axons, and synapses to meet high bioenergetic demands via fission and fusion events (13). RGCs use approximately 90% of mitochondrial-generated ATP to maintain membrane dynamics essential to sustain action potentials and survival (14–17). In addition to providing the majority of cellular ATP energy via oxidative phosphorylation (OXPHOS), mitochondria also play a vital role in RGCs by regulating Ca2+ homeostasis, apoptotic and redox signaling, and promoting neurotransmission and synaptic plasticity (18, 19). Thus, mitochondrial dysfunction in RGCs can profoundly affect neuronal function and survival. Emerging evidence of mitochondrial dysfunction has been identified in a wide variety of age-related and neurodegenerative diseases, including glaucoma (15, 16, 20–30). In a recent clinical trial, a spectrum of mitochondrial abnormalities, including increased mitochondrial DNA (mtDNA) mutations and reduced mitochondrial respiratory functions, has been reported in patients with primary open-angle glaucoma (POAG), the most common form of glaucoma (31). Also, exposure of cultured RGCs to elevated hydrostatic pressure has been shown to alter mitochondrial dynamics with reduced ATP production (26, 32). However, there is no definitive evidence of the underlying mechanisms contributing to mitochondrial dysfunction and glaucomatous neurodegeneration.

Mitochondria within RGCs are in a vulnerable condition due to RGC’s postmitotic state, high metabolic demands, increased oxidative stress, challenges in mitochondrial transport, and exposure to environmental and pathological stressors (16, 33–35). As a result, the stressed or dysfunctional mitochondria within RGCs undergo constant repair or degradation. Consequently, the cellular processes regulating mitochondrial turnover are essential for the survival and proper functioning of RGCs. Autophagy is an intracellular degradation process of damaged organelles and proteins via lysosomes. Autophagic degradation of damaged mitochondria is termed mitophagy, a major mitochondrial quality control mechanism in neurons that allows selective degradation of damaged mitochondria (36–38). In RGCs, PTEN-induced kinase 1 (PINK1)/Parkin-mediated mitophagy is the major pathway that eliminates damaged mitochondria (39). In response to stress or damage, PINK1 gets stabilized on the outer mitochondrial membrane (OMM) due to the loss of mitochondrial membrane potential (ΔΨm) and promotes recruitment of Parkin. Parkin, an E3 ubiquitin (Ub) ligase, ubiquitinates several OMM components, and the poly-Ub chains are subsequently phosphorylated by PINK1. Adaptor proteins (including p62 and optineurin (OPTN)) recognize the phosphorylated poly-Ub chains on mitochondrial proteins and initiate autophagosome formation through binding with LC3 and promote the clearance of damaged mitochondria via lysosomal degradation (40, 41). Importantly, genetic studies using a large cohort of patients have identified mutations in mitophagy-related genes, such as TNK- binding kinase 1 (TBK1) and OPTN, associated with glaucomatous neurodegeneration (27, 42–44). Furthermore, polymorphisms in the OPTN gene are associated with open-angle glaucoma, even when the IOP is normal (45), highlighting the importance of mitophagy in neuronal homeostasis and survival. The role of mitophagy/autophagy in glaucomatous neurodegeneration has been investigated by several independent laboratories (22, 23, 34, 46–56). These studies clearly demonstrated that mitophagy/autophagy is dysregulated in RGCs in response to IOP elevation. However, autophagy has been shown to play a dual role in neuroprotection and neuronal death depending on experimental conditions. These discrepancies are likely due to differences in animal models used to study glaucoma, non-specific modulators of mitophagy/autophagy, and differences in the duration of pathology and species.

To understand the role of mitophagy and mitochondrial dysfunction in the etiology of glaucomatous neurodegeneration, we have utilized two different mouse models of glaucoma, including glucocorticoid (GC)-induced OHT (57, 58) and Cre-inducible mutant myocilin (Tg. Cre-MYOC^Y437H^) mouse model of POAG (59, 60). Both mouse models closely mimic the clinical and morphological characteristics of human POAG, including IOP elevation due to TM dysfunction leading to well-defined early events of glaucomatous neurodegeneration. Using these two mouse models of glaucoma and age-matched human retinal tissue from normal and glaucoma donors, we demonstrate that IOP elevation leads to the accumulation of dysfunctional mitochondria in RGCs, ON, and ONH. Using transgenic Mt-Keima mitophagy reporter mice and RGC-specific conditional knock-out mice of autophagy, we further show that defective mitophagy is a key contributing factor for dysfunctional mitochondrial accumulation and glaucomatous neurodegeneration. Interestingly, enhancing mitophagy using either the pharmacological agent Torin 2 or RGC-specific overexpression of the Parkin gene completely rescued glaucomatous neurodegeneration in a mouse model of glaucoma and *ex vivo* cultured human retinal explants.

## Results

### Ocular hypertension leads to the accumulation of dysfunctional mitochondria in RGCs and ON axons in mouse models of glaucoma

To investigate whether mitochondrial dysfunction is associated with glaucomatous neurodegeneration, we have utilized a mouse model of GC-induced glaucoma and a recently developed Cre-inducible mouse model of myocilin POAG (Tg.Cre-MYOC^Y437H^). Both models mimic human POAG clinically and morphologically, including IOP elevation due to TM dysfunction, structural and functional loss of RGCs, and ON degeneration (57, 59). GC-induced OHT was achieved by administrating dexamethasone (Dex) via the periocular route (200 µg/eye) once weekly for 10 weeks, as described previously (57, 58, 61, 62). Dex-injected eyes showed significant IOP elevation compared to the Veh-injected control eyes (**Figure 1a)**. Consistent with our previous study (57), Dex-induced IOP elevation for 10 weeks resulted in significant loss of RGCs (**Figure 1b**) and ON axonal degeneration (**Figure S1a**). We next utilized this mouse model to evaluate whether Dex-induced OHT leads to mitochondrial dysfunction in RGCs. Transmission Electronic Microscopy (TEM) on cross sections of ON revealed that glaucomatous neurodegeneration is associated with thinning of the myelin sheath, glial scar formation, and accumulation/an increased number of mitochondria within the degenerating axons of Dex-induced OHT eyes compared to the Veh-injected eyes (**Figure 1c**). Notably, higher magnification TEM images indicated that the accumulated mitochondria in ON of Dex-induced OHT eyes are damaged or dysfunctional, as evident from their larger size and disorganized or fragmented cristae (**Figure 1d**). We further confirmed these findings by immunostaining mitochondrial marker TOM20 and oxidative DNA damage indicator 8-Hydroxy-2’- deoxyguanosine (8-OHdG) (which detects both nuclear and mitochondrial DNA damage) on retinal cross-sections. Consistent with TEM analysis, we observed increased TOM20 (**Figures S1b and S1c**) and 8-OHdG staining (**Figures 1e and S1d**) in the RGC layer (RGCL) of Dex-induced OHT eyes compared to the Veh-injected control eyes. These data indicate that chronic IOP elevation leads to the accumulation of damaged mitochondria in RGCs and oxidative DNA damage to RGCs in a mouse model of Dex-induced glaucoma.

**Figure 1:**
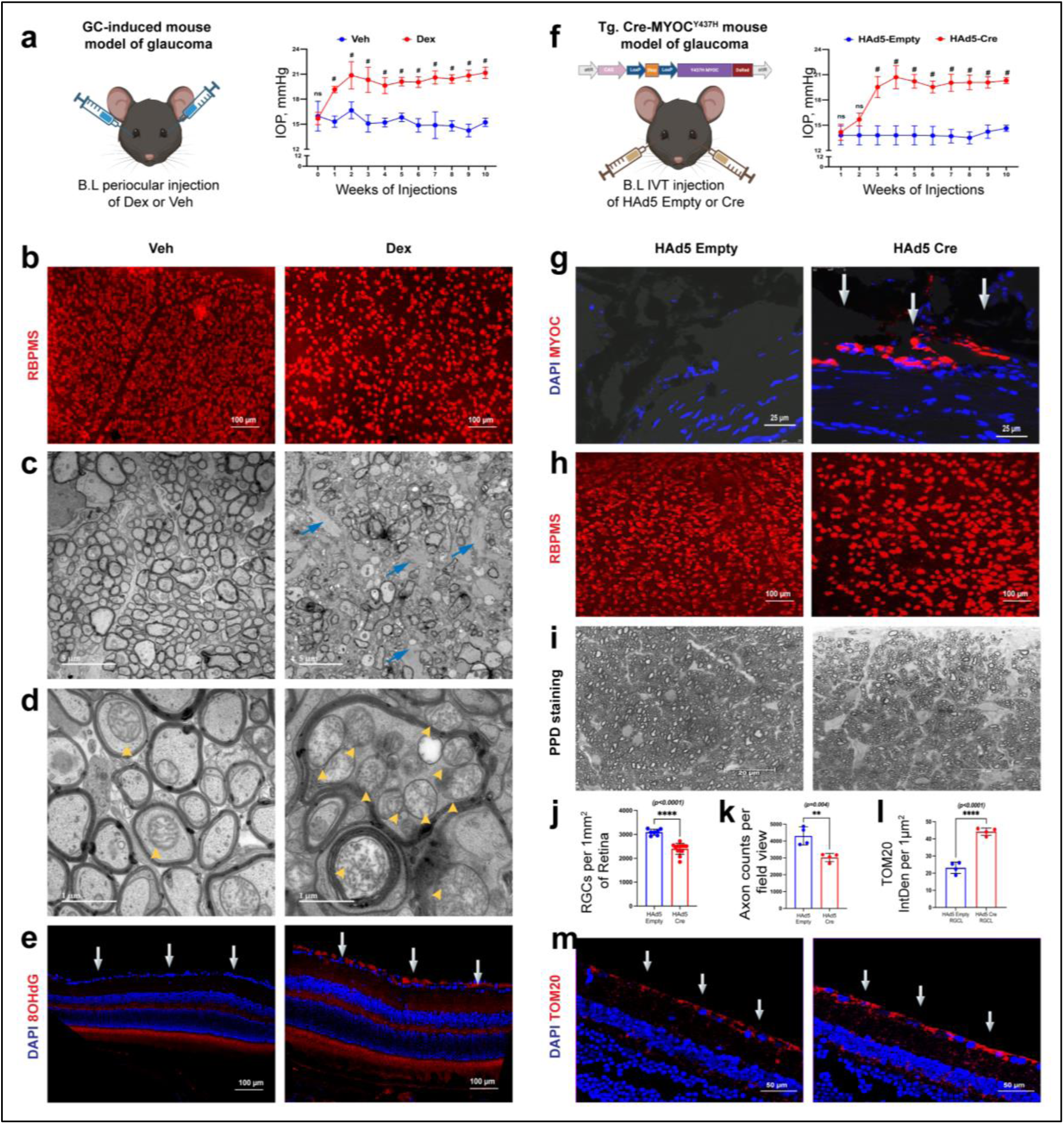
IOP elevation leads to the accumulation of dysfunctional mitochondria in RGCs in mouse models of glaucoma. **a)** Three-month-old C57BL/6J mice received bilateral (B.L) periocular injections of Veh or Dex (200 μg/eye) once a week for 10 weeks. Dex-injected mice showed significant IOP elevation starting from the first week of injections compared to Veh-injected mice (n = 10 in each group, 2-WAY ANOVA with multiple comparisons, *#p < 0.0001*, ns-not significant). **b)** Chronic elevation of IOP in Dex-injected mice resulted in a significant loss of RGCs (n=4 in each group) based on the whole mount retinal immunostaining with RGC-specific marker, RBPMS. TEM imaging of ON cross sections showed **c)** severe axonal degeneration in Dex-injected eyes compared to Veh-injected eyes (blue arrows represent glial scar), and **d)** the accumulation of dysfunctional mitochondria in Dex-induced OHT eyes compared to Veh-injected eyes (yellow arrowheads represent mitochondria within the axons); n=4 in each group. Immunostaining data showed an increased expression of **e)** 8-OHdG in RGCL of Dex-induced OHT eyes compared to Veh-injected eyes (n = 3 to 4 per group; arrows represent RGCL). **f**) Three-month-old Tg.Cre-MYOC^Y437H^ mice were injected intravitreally with either HAd5 empty or Cre (2×10^7^ pfu/eye), and IOP was monitored on a weekly. A significant IOP elevation was observed in Tg.Cre-MYOC^Y437H^ mice injected with HAd5-Cre compared to HAd5-empty injected mice (n = 4 to 5 in each group, two-way ANOVA, ns-not significant*, # p <0.0001*). **g**) Imaging of anterior segment cross-sections showed the accumulation of DS-Red tagged mutant myocilin specifically in the TM region (arrows) of HAd5-Cre-injected Tg.Cre-MYOC^Y437H^ mice compared to HAd5-empty-injected mice. Chronic IOP elevation in Tg.Cre-MYOC^Y437H^ mice resulted in **h & j)** a significant loss of RGC (∼22% loss, n=7 to 13 in each group, *p<0.0001*); **i & k)** ∼30% of axonal degeneration in PPD stained ON (n=4 in each group, *p=0.004*) and **l & m)** an increased expression of TOM20 in RGCL of in HAd5-Cre-injected Tg.Cre-MYOC^Y437H^ mice compared to HAd5-empty injected mice (n=4 in each group, *p<0.0001*).

We next explored whether the accumulation of mitochondria is also associated with glaucomatous neurodegeneration in a mouse model of MYOC-associated POAG. We recently developed a Cre-inducible transgenic mouse expressing a DsRed-tagged Y437H mutant of human myocilin (Tg.Cre-MYOC^Y437H^). Upon introduction of Cre, the Stop cassette, which inhibits transgene expression, is excised, and mutant MYOC-DsRed is expressed specifically in the TM tissue, elevating IOP in Tg.Cre-MYOC^Y437H^ mice (59, 60). Adult Tg.Cre-MYOC^Y437H^ mice were injected intravitreally with helper adenovirus (HAd) 5 expressing empty cassette or Cre (2×10^7^ pfu/eye). Expression of mutant MYOC in the TM resulted in sustained and significant IOP elevation starting at 2 weeks of injection (**Figures 1f and** **1g**). Chronic IOP elevation for 10 weeks led to glaucomatous neurodegeneration, including significant loss of RGC (**Figures 1h and** **1j**) and ON axonal degeneration (**Figures 1i and** **1k**). Similar to the GC-induced mouse model of glaucoma, glaucomatous neurodegeneration in Tg.Cre-MYOC^Y437H^ mouse model was also associated with the accumulation of mitochondria in RGCL (**Figures 1l and** **1m**) and degenerative axons (**Figure S1e)**. These data indicate that glaucomatous neurodegeneration is associated with the accumulation of dysfunctional mitochondria in RGCs.

### Ocular hypertension impairs mitophagy in RGCs, leading to the accumulation of dysfunctional mitochondria prior to glaucomatous neurodegeneration

We next hypothesized that sustained OHT leads to impaired mitophagy, resulting in abnormal accumulation of damaged mitochondria preceding neurodegeneration. We utilized the Mt-Keima reporter transgenic mouse model, which allows the measurement of mitophagy flux *in vivo*. Mt-Keima mice express a mitochondrial-targeted fluorescent reporter protein called keima, which exhibits both pH-dependent excitation and resistance to lysosomal proteases (63). The Mt-Keima protein appears red in an acidic environment (lysosomes) and green in a neutral pH environment (cytosol). Mitophagy flux is determined as mitochondria are being transferred from an autophagosome (green color) to a lysosome (red color). As shown in the schematic diagram (**Figure 2a**), Mt-Keima mice were given periocular injections of Dex (200 µg) in one eye, and the contralateral eyes were injected with Veh once a week for 5 or 10 weeks. Dex-treated eyes elevated IOP significantly compared to the contralateral Veh-injected eyes (**Figure 2b**). *In vivo* mitophagy flux was examined in 5 weeks (pre-axonal degeneration) and 10 weeks (during axonal degeneration) Veh and Dex-treated ON by confocal microscopy using excitations at 561 (red) and 458 nm (green). Representative images (**Figure 2c**) and their analyses (**Figure 2d)** demonstrated reduced red fluorescence (mitochondria in lysosomes) and increased green fluorescence (mitochondria in cytoplasm) after 5-weeks of Dex-induced OHT, indicating that mitophagy flux is significantly reduced prior to axonal degeneration. Significantly reduced mitophagy flux was observed at 10 weeks of OHT (**Figure 2e),** indicating that reduced mitophagy persists during axonal degeneration. These data indicate that mitophagy impairment precedes glaucomatous neurodegeneration. To further confirm these findings, we examined the levels of the mitophagy cargo marker p62 (SQSTM1) in mouse models of glaucoma. An increased expression of p62 protein was observed in RGCs of GC-induced and Tg.Cre-MYOC^Y437H^ mouse models of glaucoma (**Figures S2 and 2f).** These findings indicate that impaired mitophagy is a key contributing factor to the accumulation of dysfunctional mitochondria and glaucomatous progression.

**Figure 2:**
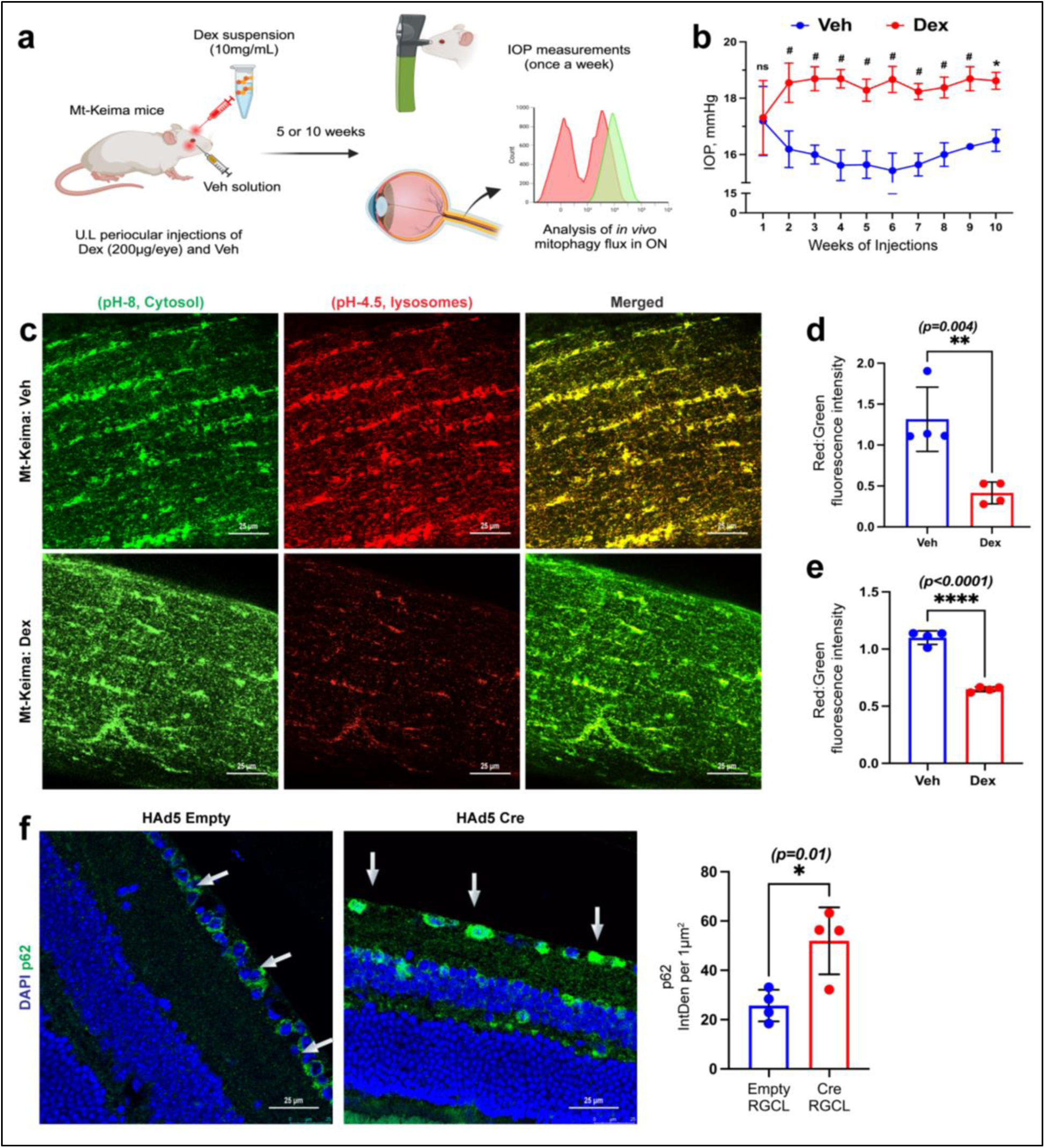
Mitophagy impairment precedes glaucomatous neurodegeneration. **a)** A schematic illustration of the study design. Mt-Keima mitophagy reporter mice were injected unilaterally with either Dex (200 μg/eye) or Veh once a week for 5 or 10 weeks via the periocular route. IOPs were recorded weekly. Following 5- and 10-weeks of OHT, the mitophagy flux was measured in the ON of Mt-Keima mice treated with Veh or Dex. **b)** Dex-treated Mt-Keima mice eyes demonstrated sustained and significant IOP elevation compared to contralateral Veh-injected eyes (n = 7 in each group, 2-WAY ANOVA with multiple comparisons, #*p* < 0.0001, ns-not significant). **c**) The representative images showing the red (excitation 561nm) and the green (excitation 458nm) fluorescence intensities in Mt-Keima mice ON after 5 weeks of Dex-induced OHT. Quantitative analysis of red: green fluorescence showed a significant reduction in mitophagy flux in **d)** 5-weeks and **e)** 10-weeks Dex-injected Mt-Keima mice ON compared to the contralateral Veh-injected ON (n=4 in each group, two-tailed unpaired t-test, \*\**p*=0.004 and \*\*\*\**p*<0.0001). **f)** Immunostaining data showed an increased expression of p62 in RGCL of HAd-Cre-injected Tg.Cre-MYOC^Y437H^ mice compared to HAd5-empty injected Tg.Cre-MYOC^Y437H^ littermates (n = 4 in each group, two-tailed unpaired t-test, \**p*=0.01 and \*\*\*\**p*=0.0001 and arrows represent RGCL).

### Mitophagy impairment and mitochondrial dysfunction in RGCs of human glaucoma donor eyes

We next examined whether impaired mitophagy and mitochondrial dysfunction are associated with glaucomatous neurodegeneration using post-mortem human donor eyes. Age-matched human normal and glaucoma donor retinas and ONs were immunostained with mitophagy markers, PINK1 and Parkin. The co-localization of these markers, observed as yellow puncta, was considered indicative of functional mitophagy. Optic nerves from glaucoma donors showed a significantly reduced number of mitophagy puncta (yellow) compared to normal eyes (**Figures 3a and** **3b**). This data strongly suggests that basal mitophagy is significantly reduced in RGCs of human glaucoma. We further quantified p62 and LC3 in the retina of these donor eyes. An increased level of p62 was observed in RGCL of glaucoma donor eyes compared to normal donor eyes, further supporting deficient mitophagy in RGCs of human glaucoma eyes **(Figures 3c and** **3d**). We also observed a significantly increased mitochondrial marker CoxIV (**Figures 3e, 3f, and S3a)**, mitochondrial oxidative stress marker SOD2 (**Figures 3g, 3h, and S3b**), and oxidative DNA damage marker (8-OHdG) (**Figures 3i, 3j and S3c**) in both RGCL and ON of the same glaucoma eyes. Together, these data indicate that impaired mitophagy, accumulation of dysfunctional mitochondria, and oxidative stress are associated with neurodegeneration in human glaucoma.

**Figure 3:**
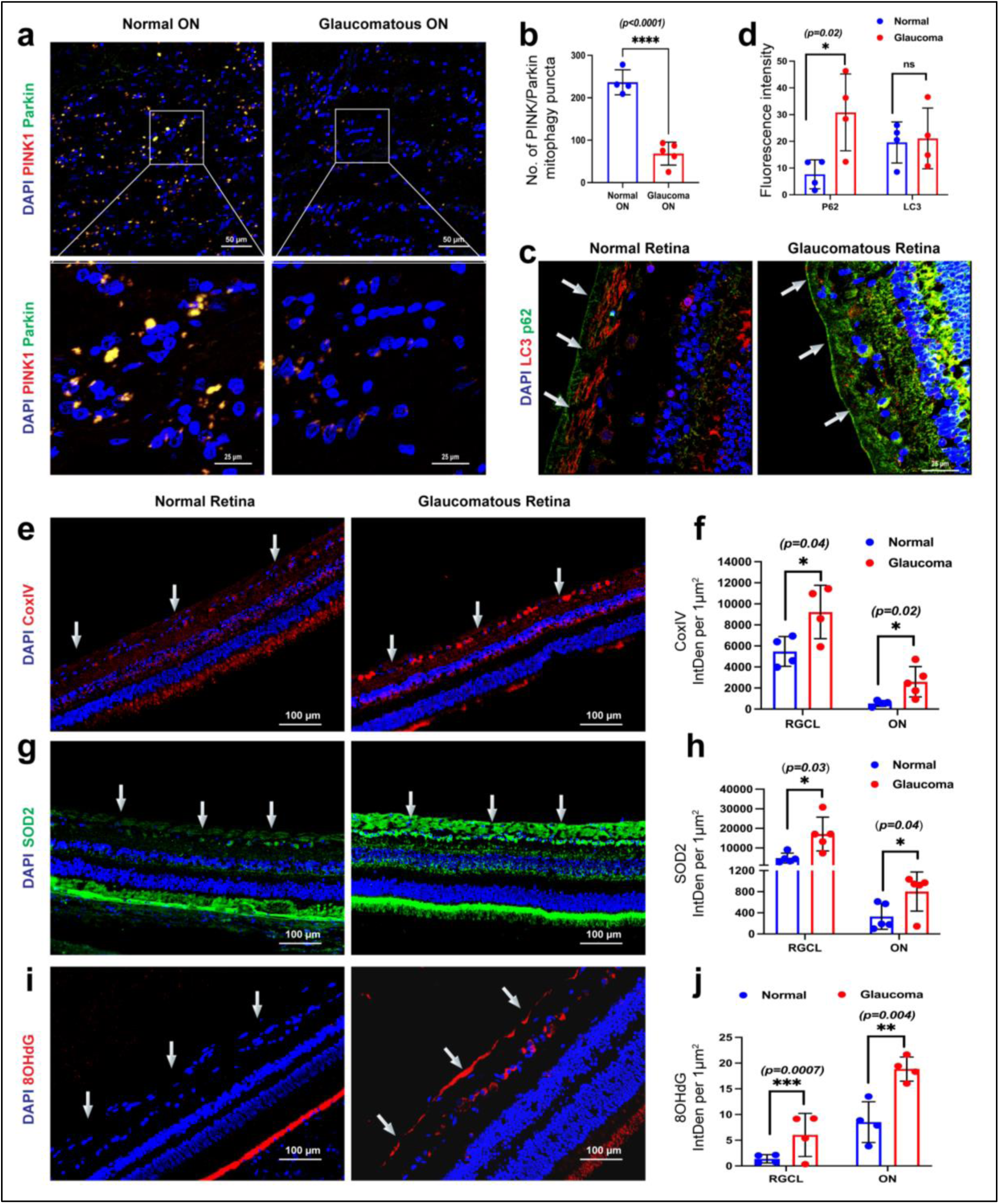
Impaired mitophagy and mitochondrial dysfunction are associated with human glaucomatous neurodegeneration. Age-matched normal and glaucomatous human postmortem retinas and ON tissues were immunostained for mitophagy and mitochondrial markers, and their fluorescence intensities were analyzed by image J. **a & b)**. Co-immunostaining of PINK1 and Parkin and their analysis demonstrated significantly decreased PINK1 and Parkin colocalization (yellow puncta) in glaucomatous ON compared to normal ON. (n = 4 normal and 5 glaucoma eyes, two-tailed unpaired t-test, *p*<0.0001). **c & d)** p62 and LC3B immunostaining, along with their analysis, demonstrated a significant increase in p62 in glaucomatous RGCL (n = 4 per group, two-tailed unpaired t-test, *p=0.02*). An increased accumulation of mitochondrial markers CoxIV (**e & f)**, SOD2 (**g & h),** and 8-OHdG (**i & j**) was observed in glaucomatous RGCL and ON compared to retinas from normal donors (n=4 to 5 in each group, two-tailed unpaired t-test). Arrows represent RGCL.

### Loss of autophagy in RGCs leads to the accumulation of dysfunctional mitochondria and subsequent neurodegeneration

Since autophagy machinery is required for functional mitophagy, we sought to determine whether conditional loss of autophagy selectively in RGCs leads to the accumulation of dysfunctional mitochondria and subsequent neurodegeneration using Atg5*^flox/flox^* mice. *The Atg5* gene encodes ATG (autophagy-related gene) 5 proteins, which serve a critical role in autophagy as a part of the ATG12-ATG5-ATG16L1 complex. ATG5 interacts with mitochondrial receptors or adaptors such as PINK1 and Parkin and facilitates the recognition, sequestration, and degradation of damaged mitochondria via the lysosomal degradation pathway (64, 65). Conditional knockdown of *Atg*5 selectively in RGCs was achieved by intravitreal injection of adeno-associated viral vector (AAV) 2 expressing Cre-recombinase under the control of synapsin promoter (AAV2-Syn-Cre). 3-month-old Atg5*^flox/flox^* mice were injected intravitreally with AAV2-Syn-Cre (1.67E+13 VG/mL) in one eye, and the contralateral eyes were injected with AAV2- Syn-Empty. Structural and functional loss of RGCs and axonal degeneration were examined at 6 weeks after the injection (**Figure 4a**). To first determine whether AAV-Syn exhibits specific tropism to RGCs, C57BL/J mice were injected with AAV-Syn-GFP. Retinal cross-sections (left panel) and whole mount retina (right panel) staining with RGC marker, RBPMS revealed selective GFP expression in RBPMS-positive RGCs, indicating that AAV-Syn exhibits specific tropism to RGCs (**Figure 4b**). Loss of RGC-specific *Atg*-5 resulted in the accumulation of p62 and CoxIV mitochondrial markers in RGCL with increased 8-OHdG levels in ON (**Figures 4c and** **4d**). Furthermore, TEM images confirmed the accumulation of damaged or dysfunctional mitochondria in ON of *Atg*5 knock-out eyes (**Figure 4e**). These data indicate that impaired mitophagy due to RGC-specific loss of *Atg5* leads to the accumulation of mitochondria and oxidative DNA damage in RGCs. Next, we examined whether loss of autophagy in RGCs is sufficient to induce neurodegeneration in mice. PERG analysis demonstrated a significant functional loss of RGCs in Atg5*^flox/flox^* mice eyes injected with AAV2-Syn-Cre (**Figures 4f and** **4g**). The knockdown of RGC-specific *Atg*5 resulted in a significant loss of RGC (∼39%) (**Figures 4h and i**) and severe axonal loss (∼59%) after 6 weeks of injection (**Figures 4j and** **4k**). Together, these data indicate that loss of RGC-specific autophagy leads to abnormal accumulation of dysfunctional mitochondria in RGCs, leading to neurodegeneration.

**Figure 4:**
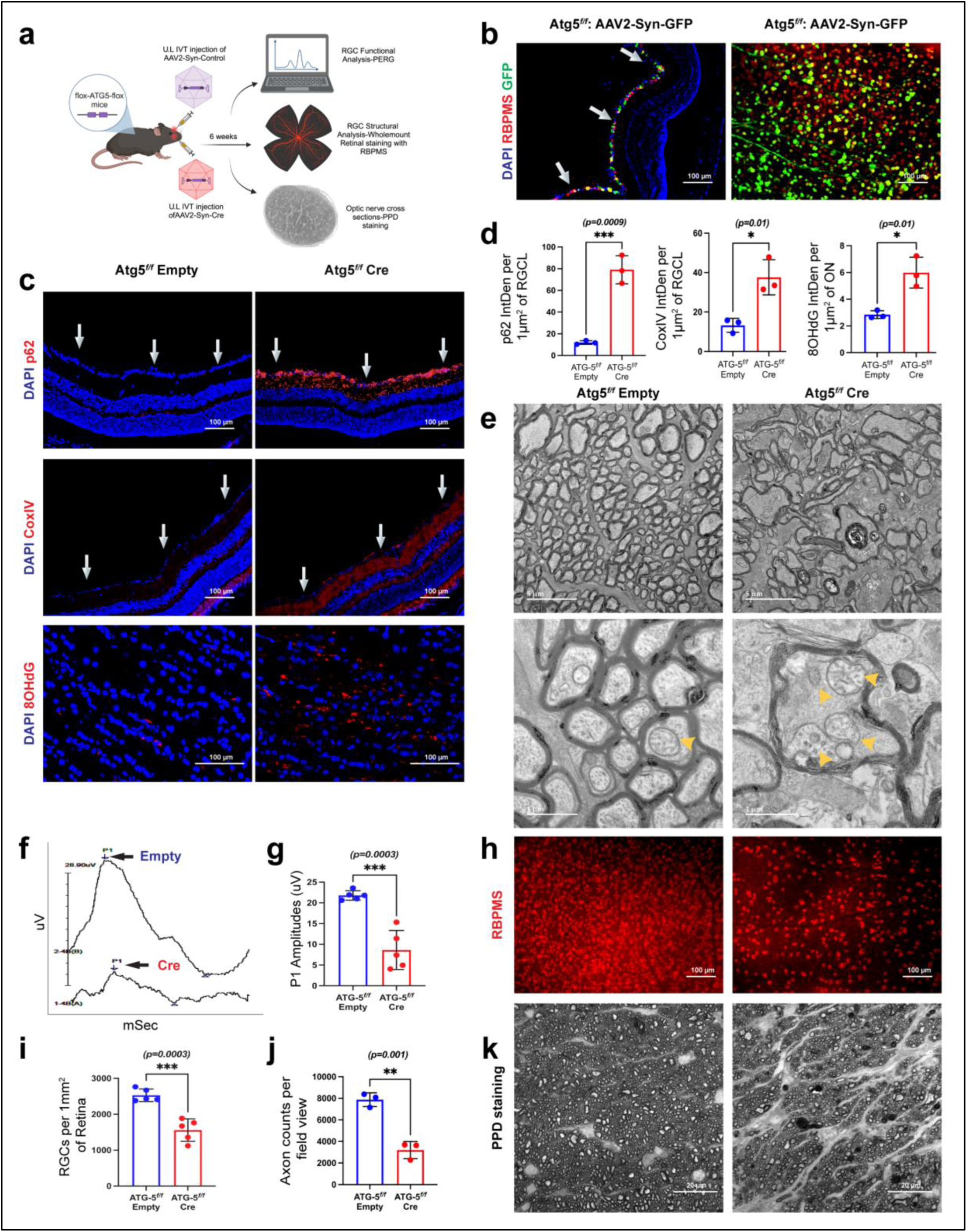
RGC-specific conditional knockdown of *Atg*5 induces the accumulation of dysfunctional mitochondria, leading to neurodegeneration. **a)** A schematic illustration of the study design. Three-month Atg5*^flox/flox^* mice were intravitreally injected in one eye with AAV2-Syn-Cre and the contralateral eye with an AAV2-Syn-empty vector. Mitochondrial accumulation and neurodegeneration were evaluated after 6 weeks of injection. **b)** AAV2-Syn-GFP was injected intravitreally, and GFP expression in RGC was examined in the retinal cross-section (left panel) and whole mount retina (right panel). Co-labeling of GFP with RBPMS demonstrated a specific tropism of AAV2-Syn-GFP to RGCs. **c & d**) Representative Immunostaining images and their intensity measurements demonstrate significantly increased levels of p62 and CoxIV in RGCL along with 8-OHdG in ON of AAV2-Syn-Cre injected eyes compared to contralateral AAV2-Syn-empty vector injected eyes of Atg5*^flox/flox^* mice (Arrows represent RGCL, (n=3 in each group, two-tailed unpaired t-test). **e**) Representative TEM images show accumulation of dysfunctional mitochondria in AAV2-Syn-Cre injected eyes compared to empty vector injected eyes of Atg5*^flox/flox^* mice (yellow arrowheads represent mitochondria within ON axons); n=4 in each group. RGC- specific conditional knockdown of the *Atg*5 resulted in (**f** & **g**) decreased RGC function as evident from reduced pattern ERG amplitudes (21.78μV vs 8.61μV) (n=5 in each group), (**h & i**) a significant RGC loss, as determined by whole mount retinal staining with RBPMS (∼39% loss) (n=6 in each group, and (**j & k**) a significant loss of healthy axons (∼59%), as determined by PPD staining 6 weeks after AAV2-Syn-Cre injection. (n=3 in each group, two-tailed unpaired t-test).

### Enhancement of mitophagy via Torin 2 prevents glaucomatous neurodegeneration in a mouse model of glaucoma and *ex vivo* human retinal explant model

Since our results indicate impaired mitophagy as a key player in glaucomatous neurodegeneration, we explored whether enhancing mitophagy in RGCs would ameliorate glaucomatous neurodegeneration. We examined whether Torin 2, which is a potent and selective inhibitor of the mammalian target of rapamycin (mTOR) kinase, enhances mitophagy and rescues a mouse model of glaucoma. First, we investigated whether Torin 2 enhances mitophagy flux using Mt-Keima mice. 3-month-old Mt-Keima mice were intravitreally injected with Torin 2 (1 mM dissolved in DMSO) in one eye, and the contralateral eyes were injected with DMSO as a control. I*n vivo* mitophagy flux measurements demonstrated a significant enhancement of mitophagy flux 24 hrs after Torin 2 treatment (**Figures 5a & b**). Next, we examined whether Torin 2 prevents neurodegeneration in a mouse model of GC-induced glaucoma. 3-month-old C57BL/6J mice were treated with Dex once a week for 7 weeks. IOPs were monitored to ensure significant IOP elevation. At 7 weeks of OHT, Dex-treated mice were injected intravitreally with Torin 2 (1 mM dissolved in DMSO) in one eye, and the contralateral eyes were injected with DMSO as a control. Dex-treatment was continued for another 3 weeks to maintain OHT **(Figure 5c)**. IOP measurements showed that Dex-treated eyes continued to have significant IOP elevation, and Torin 2 did not alter IOPs in both control and Dex-treated eyes (**Figure 5d**). At the end of the 10-week study period, PERG measurements revealed that Torin 2 treatment significantly improved RGC function in Dex-treated (Dex^Torin2^) eyes, as evident from significantly higher PERG amplitude compared to control Dex-treated eyes (Dex^DMSO^) (**Figures 5e and** **5f**). Consistent with improved RGC function, we observed that Torin 2 treatment significantly preserved the total number of surviving RGCs and healthy axons in Dex^Torin2^ eyes compared to Dex^DMSO^ control eyes (**Figures 5g and** **5h**). Together, these data suggest that enhancement of mitophagy via Torin 2 protected glaucomatous neurodegeneration in a mouse model of Dex-induced glaucoma. We further confirmed these findings using an *ex vivo* human retinal explant model (66, 67). The retinal explants were treated with either DMSO or Torin 2 (100 μM) and cultured for 7 days in an explant medium under neurotrophic factors-deprived conditions. The retinal explants were then fixed, and surviving RGCs were quantified using an anti-RBPMS antibody (**Figure 5i**). Torin 2 significantly increased the total number of surviving RGCs compared to DMSO-treated controls (**Figure 5j**), indicating the neuroprotective role of mitophagy in human RGC survival.

**Figure 5:**
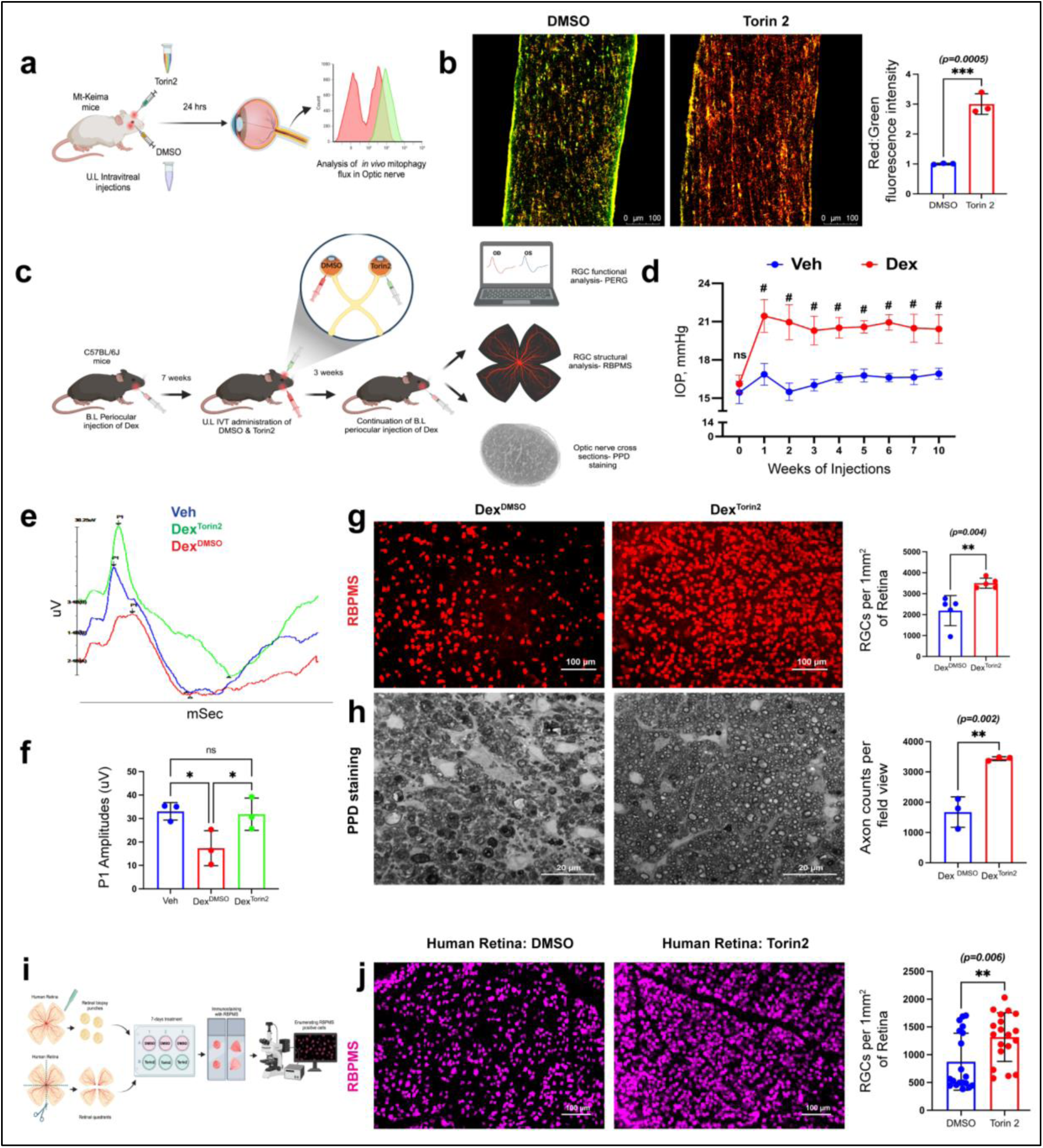
Enhancing mitophagy via Torin 2 prevents glaucomatous neurodegeneration. **a)** A schematic illustration of the study design. Mt-Keima mice were intravitreally injected with Torin 2 (1 mM) in one eye and DMSO in the contralateral eye. Mitophagy flux was examined after 24 hrs of treatment. **b)** The representative images and their fluorescence intensity analysis showed a significant increase in red-to-green ratio (mitophagy flux) in the Torin 2 treated ON compared to the contralateral DMSO treated ON of Mt-Keima mice (n=3 in each group, two-tailed unpaired t- test, \*\*\**p*=0.0005). **c)** The study design illustrating the Torin 2 treatment plan in a mouse model of GC-induced glaucoma. 7-week Dex-induced IOP elevated C57BL/6J mice were treated intravitreally with Torin 2 (1 mM) in one eye, and the contralateral eyes were injected with DMSO control. A sustained elevation of IOP was maintained for another three weeks by administering Dex periocular injections. After 10 weeks of injection, glaucomatous neurodegeneration was assessed. **d**) Dex-injected mice demonstrated a significant and sustained IOP elevation throughout the course of the study. #*p*<0.0001, n=6 to 10 eyes in each group; two-way ANOVA with multiple analysis. **e & f)** The representative PERG waves and statistical analysis showed that Torin 2 significantly improved PERG amplitudes in Dex-treated eyes compared to Dex and DMSO-treated eyes Dex (n = 3 in each group, two-tailed unpaired t-test). **g**) The representative whole mount retinal staining with RBPMS (n=5 in each group) and **h)** PPD stained ON axons (n=3 in each group) showed an increased number of RGCs and healthy axons in Dex-induced OHT eyes treated with Torin 2 compared to contralateral Dex-induced OHT eyes treated with DMSO respectively (two-tailed unpaired t-test). **i**) A study design illustrating the Torin 2 treatment plan in human retinal explants. Retinal explants were cultured for 7 days under neurotrophic factors deprivation conditions and treated with DMSO or Torin 2 (100 μM) in an explant medium and quantified surviving RGCs using an anti-RBPMS antibody. **j)** Torin 2 treated retinal explants showed a higher number of RGCs than those treated with DMSO (n=4 in each group, two-tailed unpaired t-test, *p*=0.006).

### Genetic induction of mitophagy via RGC-specific Parkin overexpression rescues glaucomatous neurodegeneration in a mouse model of glaucoma

We next investigated whether genetic induction of mitophagy by expressing Parkin in RGCs rescues glaucomatous neurodegeneration in a mouse model of Dex-induced glaucoma (**Figure 6a**). Three-month-old C57BL/6J mice were intravitreally injected with either AAV2-Syn-Parkin or AAV2-Syn-empty. IOP was elevated using periocular injections of Dex once a week for 10 weeks. We noticed that Parkin overexpression did not affect IOPs significantly in Dex-injected mice (**Figure 6b**). Overexpression of Parkin significantly reduced CoxIV and p62 levels in Dex^Parkin^ eyes compared to Dex^Empty^ eyes (**Figures 6c and** **6d**). TEM imaging further confirmed that RGC-specific Parkin overexpression prevented the accumulation of dysfunctional mitochondria and restored a healthy mitochondrial pool in ON axons (**Figure 6e).** Importantly, overexpression of Parkin in Dex^Parkin^ mice exhibited improved PERG amplitudes (**Figure 6f)** and showed significantly higher total RGCs (**Figures 6g and** **6h**) and healthy axons compared to Dex^Empty^ mice group (**Figures 6i & j**). Together, these data strongly suggest that the enhancement of mitophagy by overexpression of Parkin rescues glaucomatous neurodegeneration in a mouse model of glaucoma.

**Figure 6:**
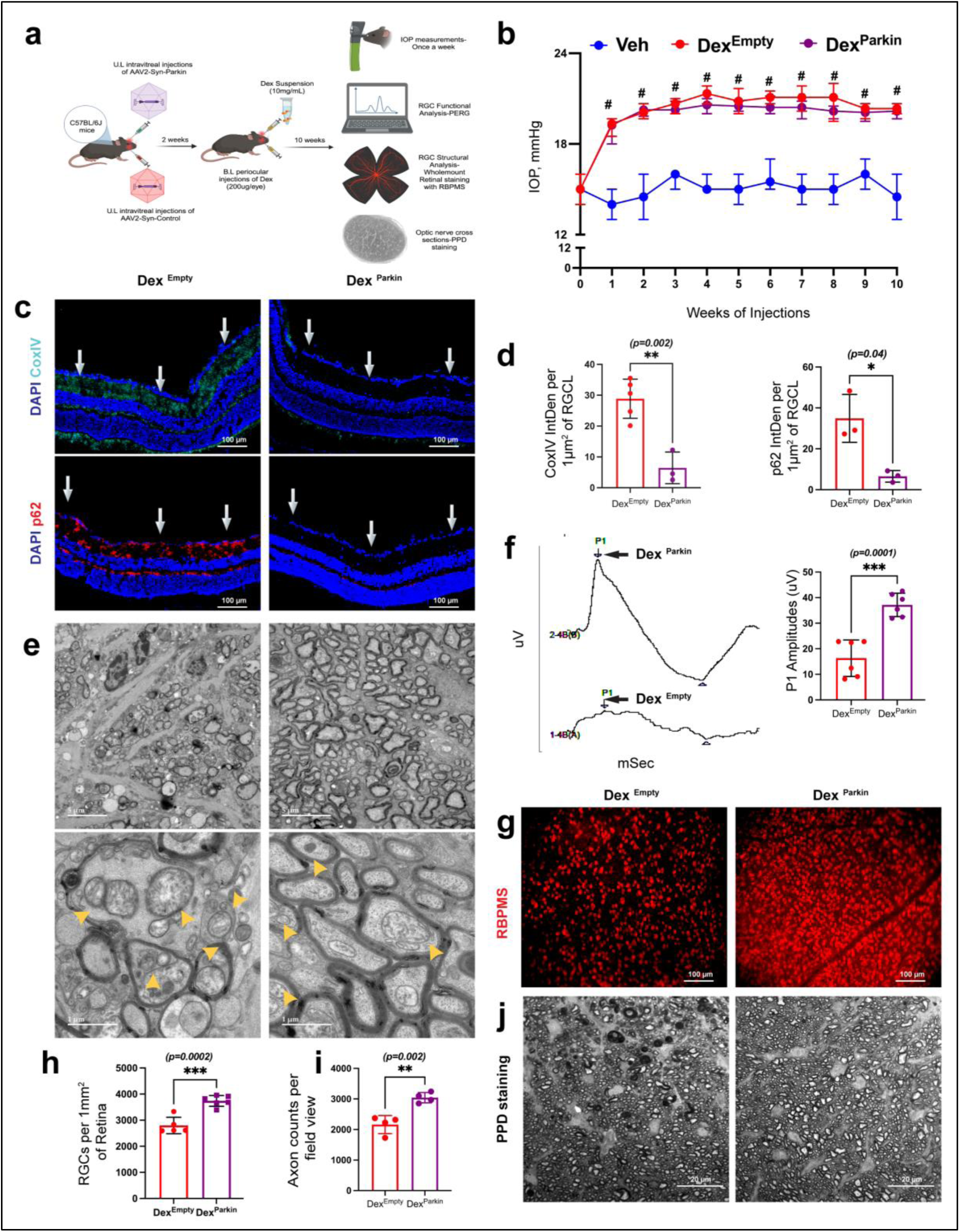
RGC-specific Parkin overexpression rescues glaucomatous neurodegeneration in a mouse model of glaucoma. **a)** A schematic illustration of the study design of Parkin overexpression in a mouse model of Dex-induced glaucoma. Three-month-old C57BL/6J mice were intravitreally injected with either AAV2-Syn-Parkin or AAV2-Syn-empty. After 2 weeks of injection, IOP was elevated bilaterally via periocular injections of Dex once a week for 10 weeks, and glaucoma phenotypes were examined. **b**) Dex-treated OHT mice (Dex^Parkin^ and Dex^Empty^) showed a significantly higher IOP than Veh-injected mice and no significant differences in IOPs were observed between Parkin overexpressed (Dex^Parkin^) and empty vector (Dex^Empty^) Dex-treated mice (n = 10 to18 in each group, 2-WAY ANOVA with multiple comparison, #*p < 0.0001*, ns-not significant). **c & d**) Immunostaining and its analyses revealed a reduced expression of CoxIV and p62 in Dex^Parkin^ eyes compared to Dex^Empty^ eyes (n = 3 to 5 in each group). Overexpression of Parkin in RGCs exhibited significantly improved **e**) healthy mitochondrial pool in ON as evident from representative TEM images, **f**) PERG amplitudes, **g & h**) surviving RGC, and **i & j**) healthy axons in PPD stained ON of Dex^Parkin^ eyes compared to Dex^Empty^ eyes (n=4 to 6 in each group, two-tailed unpaired t-test).

## Discussion

Although the role of mitochondrial dysfunction in glaucomatous neurodegeneration is well-recognized, the underlying mechanisms of increased accumulation of dysfunctional mitochondria in RGCs are poorly understood. Here, we show that mitophagy is impaired in RGCs of mouse models of glaucoma and human glaucoma donor eyes. Importantly, OHT impairs mitophagy, resulting in the accumulation of dysfunctional mitochondria in RGC soma and axons, which leads to chronic oxidative stress, DNA damage, and subsequent neurodegeneration. Consistent with this observation, we further show that basal autophagy/mitophagy in RGCs is required to maintain healthy mitochondria and neuronal survival. Notably, this is the first study demonstrating that impaired mitophagy and mitochondrial dysfunction precede glaucomatous neurodegeneration. Importantly, genetic or pharmacological activation of mitophagy ameliorated glaucomatous neurodegeneration in mouse models of glaucoma and *ex vivo* human retinal explant model.

Several independent laboratories have demonstrated that altered mitochondrial dynamics and metabolic stress due to mitochondrial dysfunction are associated with glaucomatous neurodegeneration (68–72). Recent studies have shown that mitochondrial dysfunction leads to metabolic perturbations, and dietary supplements with nicotinamide and pyruvate enhance neuronal function in open-angle glaucoma patients (73–75), suggesting the importance of mitochondria in neuronal function. Despite the well-known disease relevance of mitochondrial dysfunction in glaucoma, the underlying molecular mechanisms are poorly understood. Mitophagy, a highly conserved cellular process, plays a pivotal role in mitochondrial quality control by sequestering defective mitochondria into mitophagosomes for subsequent lysosomal degradation, maintaining mitochondrial and metabolic balance, energy supply, neuronal survival, and health (76, 77). On the other hand, impaired mitophagy promotes the accumulation of damaged mitochondria, leading to cellular dysfunction and neurodegeneration. There is mounting evidence from *in vitro* and *in vivo* studies that mitophagy/autophagy has neuroprotective effects on RGCs in ocular neurodegenerative diseases, including glaucoma (78–81). Using Morrison’s model of OHT, we have recently demonstrated changes in mitophagosome formation that predispose RGCs to neurodegeneration in rats (82). In contrast, some studies have suggested that activation of mitophagy/autophagy may contribute to the death of RGCs under glaucomatous conditions (48, 55, 56, 83, 84). These discrepancies in findings are likely due to animal models utilized to study the role of autophagy/mitophagy in RGCs. Other concerns with these previous studies are the timings of mitophagy/autophagy analyses during glaucoma pathology and the use of non-selective drugs to induce mitophagy or autophagy. Furthermore, most of these studies focused only on global autophagy, ignoring mitochondrial morphological changes and their impact on oxidative stress, one of the driving factors for glaucoma progression. In this study, we have used mouse glaucoma models that replicate human glaucoma pathology. Both GC and mutant MYOC-induced OHT lead to glaucomatous neurodegeneration similar to human glaucoma, making these two mouse models ideal for studying the etiology of human glaucoma. In addition, well-defined timeframes of neurodegeneration in these mouse models allow us to study the exact timings of RGC pathology. Importantly, we have investigated the role of mitophagy in both RGC soma and axons using different mouse models, including GC and mutant MYOC-induced mouse models of glaucoma, mitophagy reporter transgenic Mt-Keima mice, and autophagy-specific Atg5*^f/f^* mice. Importantly, in contrast to existing studies, we have utilized age-matched human postmortem donor tissue to correlate our findings from mouse models. Our studies also utilized pharmacological and genetic approaches to activate autophagy/mitophagy in RGCs to rescue glaucoma.

PINK1-Parkin-mediated mitophagy is a major and well-studied pathway in neurons, including RGCs (85–87). In this study, we utilized mitophagy reporter Mt-Keima mice, which detect PINK1- Parkin mitophagy with greater sensitivity compared to the other reporter mice (88). Using Mt-Keima mitophagy reporter mice, we analyzed the effect of chronic IOP elevation on mitophagy flux both prior to axonal degeneration (5 weeks of OHT) and during axonal degeneration (10 weeks of OHT). In our previous studies, we have shown that most of the active axonal transport persists during the early stages of ON degeneration, while complete transport deficits occur at later/severe stages of axonal damage in a mouse model of GC-induced glaucoma (57). Interestingly, mitophagy flux was reduced prior to axonal degeneration, despite active axonal transport, suggesting that mitophagy impairment precedes structural and functional loss of RGCs. In addition, axonal transport deficits due to persistent OHT may further contribute to the accumulation of damaged mitochondria and impaired mitophagy, which is solely responsible for degrading damaged mitochondria. This ultimately leads to a decrease in mitochondrial energy production and an increase in oxidative stress, which further exacerbate neuronal damage. Some studies have demonstrated that damaged mitochondria from distal axons are transported retrogradely to the neuronal soma for lysosomal degradation (89–92). However, other studies have also found that dysfunctional mitochondria are degraded locally in distal axons via mitophagy rather than returning to the soma to prevent the spread of oxidative damage (93–96). Nicholas Marsh-Armstrong et al. have also identified the existence of transcellular mitophagy at the ONH, where large numbers of mitochondria are shed from RGCs and degraded by the lysosomes of adjoining glial cells (94, 97). Furthermore, the vulnerable or stressed axonal mitochondria exhibit reduced mitochondrial motility, preventing their return to the soma for degradation (93, 98).

Our study demonstrated impaired mitophagy results in the accumulation of dysfunctional mitochondria in axons and soma of RGCs in both mouse and human glaucoma. This was evident from an increased expression of the mitochondrial marker CoxIV and the mitophagy cargo receptor p62. Moreover, TEM imaging further confirmed the accumulation of dysfunctional mitochondria with a swollen matrix and disorganized cristae. Importantly, we have seen reduced colocalization of PINK and Parkin in human glaucomatous ON. Consistent with this observation, increased 8-OHdG, a marker of oxidative damage to DNA (including nuclear DNA and mtDNA), and mitochondrial SOD2 levels in RGCs and axons indicated the presence of chronic oxidative stress. Persistent oxidative stress/damage due to defective mitophagy may lead to activation of the intrinsic (mitochondrial) apoptotic pathway by promoting the release of pro-apoptotic factors such as cytochrome c into the cytosol, which activates caspase-9 and, subsequently, caspase-3, leading to the apoptosis of RGC soma. In addition, impaired energy production, calcium dysregulation, and induction of neuroinflammation can induce structural and functional loss of RGCs, triggering neuronal degeneration. Glaucomatous neurodegeneration is compartmentalized, and distinct molecular signals contribute to the degeneration of both axons and soma (99). Our study identifies impaired mitophagy as a common pathological event underlying both axonal degeneration and RGC soma loss. Our study demonstrates that basal autophagy/mitophagy is required for maintaining healthy mitochondria, neuronal function, and survival. This is evident from the observation that *Atg*5 loss in RGCs induces neurodegeneration, accompanied by increased mitochondrial accumulation and oxidative DNA damage. We utilized AAV2-Cre recombinase under a synapsin promotor to delete Atg5 specifically from RGC, which showed more than 90% transduction ability in our studies and RGC-specific tropism.

Since mitophagy is the fundamental mechanism for mitochondrial homeostasis, therapeutic interventions aimed at mitophagy induction may ameliorate mitochondrial dysfunction and protect the RGCs from IOP-induced damage. In this study we showed enhancing mitophagy to clear the accumulated dysfunctional mitochondria as an attractive glaucoma therapeutic strategy. We utilized Torin 2 to rescue IOP-induced impaired mitophagy in a GC-induced mouse model of glaucoma. Previously, we demonstrated that Torin 2 rescues TM cell dysfunction in a mouse model of myocilin POAG by enhancing the autophagic degradation of mutant myocilin. Torin 2- induced autophagy flux was associated with reduced mutant myocilin and p62 accumulation (100). Consistent with this previously published study (100), Torin 2 enhanced mitophagy flux within 24 hr of its administration (intravitreal) in Mt-Keima mitophagy reporter mice and reduced p62 accumulation in RGCL of OHT mice. Torin 2 is a second-generation mTOR catalytic inhibitor with higher potency than rapamycin and induces mitophagy more robustly by inhibiting both mTORC1 and 2 (101). It has been shown that Torin 2 significantly improves tolerability and, hence, is useful as a therapeutic agent against aging and neurodegenerative diseases (102). One of the major strengths of our study is to examine the therapeutic potential of mitophagy enhancement via Torin 2 under mild to moderate neurodegenerative conditions. We administered Torin 2 to mice with GC-induced glaucoma after 7 weeks of OHT and maintained the elevated IOP for an additional 3 weeks to understand the disease-modifying potential of the drug. Enhancement of mitophagy via Torin 2 completely prevented neurodegeneration in our mouse model of glaucoma. Notably, mitophagy enhancement via Torin 2 increased the number of surviving RBPMS-positive RGCs in an *ex vivo* human retinal explants model, further supporting the therapeutic potential of Torin 2 glaucoma treatment.

Since Torin 2 has a wide range of activities and lacks specificity to RGCs, we further utilized genetic activation of mitophagy selectively in RGCs by expressing the mitophagy gene Parkin in these cells. As an E3 ubiquitin ligase, Parkin is recruited to the OMM in response to altered mitochondrial ΔΨm, which facilitates mitophagy by ubiquitinating mitochondrial membrane proteins (47, 103). Several studies have shown that parkin overexpression prolongs the lifespan of adult Drosophila, (104) prevents neurodegeneration in Parkinson’s disease (105–107), and protects RGCs in experimental glaucoma (47). Hass et al. also found that the knockout of mitochondrial uncoupling protein 2 prevented RGC death in a microbead occlusion mouse model of glaucoma by promoting mitophagy (22), highlighting the neuroprotective role of mitophagy. Consistent with these studies, we show that modulating mitophagy via Parkin overexpression prevented structural and functional loss of RGCs and axons in a chronic glaucoma mouse model by reducing mitochondrial accumulation and oxidative stress. Moreover, TEM imaging and immunostaining data further confirm the clearance of dysfunctional mitochondrial accumulation in RGC axons and reduced oxidative stress in RGCs with improved PERG amplitudes.

In summary, this is the first study to demonstrate that mitophagy impairment precedes glaucomatous neurodegeneration and is associated with both mouse and human glaucoma. Importantly, augmenting mitophagy through genetic or pharmacological approaches provides an attractive therapeutic strategy for preventing vision loss in glaucoma.

## Materials and Methods

### Experimental Animals

C57BL/6J were purchased from the Jackson Laboratory. *Atg*5*^flox/flox^* mice on a pure C57BL/6J genetic background were provided by Dr. Noboru Mizushima (The University of Tokyo, Tokyo, Japan) and were received from Thomas Ferguson’s lab (Washington University School of Medicine, St. Louis, Missouri, USA). Mt-Keima mice on FVB/N genetic background were kindly provided by Dr. Toren Finkel (University of Pittsburgh). *Tg.Cre-MYOC^Y437H^* mice on a pure C57BL/6J genetic background were developed in our laboratory as described recently (59). Both male and female mice between the ages of 3 and 4 months were utilized in this study. Under the controlled conditions of temperature (21 to 26°C) and humidity (40 to 70%), mice were subjected to a 12 h light/12 h dark cycle (8:00 PM to 8:00 AM). All animals were fed standard chow ad libitum and provided with water. The number of animals used in each experiment is indicated in the corresponding figure or figure legend.

### Mouse models of glaucoma

3-month-old mice were injected with 20 μL/eye of either Vehicle (Veh) or freshly prepared dexamethasone (Dex) (i.e. 200 μg/eye) suspension via the periocular route once a week for 5 or 10 weeks under isoflurane anesthetic conditions (isoflurane (2.5%); oxygen (0.8 L/min)). IOPs were monitored weekly. To induce Tg.Cre-MYOC^Y437H^ mice, 3-month-old *Tg.CreMYOC^Y437H^* mice were intravitreally injected with HAd5 expressing empty cassette or Cre (2 ×10^7^ pfu/eye) using a Hamilton syringe fitted with a sterile 33-gauge needle under isoflurane anesthetic conditions. IOPs were monitored weekly. At the end of the study period, mice were euthanized by inhalation of CO_2_ or intracardiac perfusion followed by cervical dislocation.

### IOP Measurements

IOPs were measured between 9-11 AM using a Tonolab rebound tonometry (Colonial Medical Supply) under isoflurane anesthesia as described previously (108, 109). IOP measurements were performed once a week throughout the study period in a masked manner. At each time point, an average of six IOP readings were taken from each eye and represented as the final IOP value.

### Pattern electroretinography (PERG)

PERG measurements were performed using the Miami PERG system (Jorvec, Miami, FL) according to the manufacturer’s instructions to analyze the RGC function as described previously (57, 110). Briefly, experimental and control mice were anesthetized with ketamine/xylazine solution (100/10 mg/kg) and placed on a temperature-controlled metal base at a fixed distance (10 cm) from the LED monitors to maintain a constant body temperature (37°C). The PERG was derived simultaneously from each eye using electrodes placed subcutaneously at the snout (active), back of the head (reference), and tail (ground). An average of two consecutive repetitions was taken, and amplitudes (P1-N2) representing the RGC function were shown in the results.

### Immunostaining

Enucleated eyes from experimental and control mice were fixed in 4% paraformaldehyde for 3 hr at room temperature and then processed and embedded in paraffin. 5- micron tissue sections were cut with a microtome, and immunostaining was performed as described previously (111, 112). Briefly, tissue sections were deparaffinized in xylene and rehydrated in gradient concentrations of ethanol (100, 95, 70, and 50%), followed by antigen retrieval using citrate buffer (pH = 6). Following a 15-minute wash in 1X PBS, tissue sections were incubated with blocking buffer (containing 10% goat serum and 0.5%-2% Triton-X-100 in 1X PBS) for 2 hr, followed by overnight incubation with primary antibodies in blocking buffer at 4°C. Following 3 washes in 1X PBS, tissue sections were further incubated with appropriate secondary antibodies for 2 h at room temperature. Tissue sections were then washed again with 1X PBS and mounted with a mounting medium containing DAPI nuclear stain (Vector Labs, Inc., Burlingame, CA, USA). Images were captured by either a Keyence fluorescence microscope (Itasca, IL, USA) or a confocal microscope and analyzed using ImageJ. Tissue sections incubated without a primary antibody served as a negative control, and the relative signal intensities were subtracted from an averaged background intensity.

For immunostaining human donor eyes, age-matched normal and glaucomatous retina and ON sections were utilized (**Supplementary Table 1**). The design and conduct of the study were compliant with all relevant regulations regarding the use of human participants and with the criteria established by the Declaration of Helsinki. Briefly, post-mortem fixed human eyes were obtained from the Lions Eye Institute for Transplant & Research (Tampa, FL).

### Whole-mount retinal staining and RGC counting

The total number of surviving RGCs in experimental and control mice groups was determined by whole-mount staining with RGC-specific RBPMS antibody, as described previously (57, 110). Briefly, eyes were enucleated and fixed in 4% PFA for 12 hr at 4 °C. Eyes were rinsed with 1X PBS. Anterior segment was removed, and the whole retina was carefully separated from the posterior cup. The isolated retinas were incubated at 4 °C for 12 hr in blocking buffer (10% goat serum and 0.2% Triton X-100 in PBS). The whole retinas were then incubated with RBPMS antibody for three days at 4°C and washed 3 times in 1x PBS. The retinas were further incubated with a corresponding secondary antibody (goat anti-rabbit 568, 1:500; Invitrogen) for 2 hr at room temperature, rinsed 3 times in 1X PBS, and flat-mounted on a glass slide by cutting four equal quadrants attached to the center of the retina. A Keyence microscope was used to capture the images at a magnification of 20x. For counting RGCs, at least 12-16 non-overlapping images from the entire retina, including the periphery, mid-periphery, and center of the retina, were taken, and RBPMS-positive cells were counted using ImageJ software.

### Optic nerve axonal damage by Paraphenylenediamine (PPD) staining and TEM

ON axonal degeneration from experimental and control mice groups was examined using PPD staining. Transmission electron microscopy (TEM) was used further to examine the ultrastructural morphology of degenerative axons as described previously (57, 110). ON were incubated overnight with 1% osmium tetroxide at 4°C and then rinsed with 0.1 M phosphate buffer and 0.1 M sodium-acetate buffer, followed by dehydration in graded ethanol concentrations. ON was embedded in resin (Eponate-12; Ted Pella), and semithin sections were cut and stained with 1% PPD for 10 min after embedding. On each ON, five images were taken without overlap with a Zeiss confocal microscope, and surviving axons were counted in an area of 8100 square micrometers from each image using Image J software. TEM analysis was performed on 1 μm sections of ON from experimental and control mice groups at the Electron Microscopy Core Facility at the University of Missouri, Columbia.

### Analysis of *in vivo* mitophagy flux

ON from Mt-Keima mice were carefully removed and immediately sealed under the coverslip on a glass slide and analyzed using confocal microscopy with excitations at 561 (red) and 458 nm (green). The level of mitophagy flux was calculated by dividing the area of the “red” signal (excitation by wavelength 561 nm) by the area of the “green” signal (excitation by wavelength 458 nm), using the Fiji software package, which is based on ImageJ (National Institutes of Health).

### *Ex vivo* culture of human axotomized retinal explants

Human retinas were isolated from healthy donors within 12 hr of their death from the UNTHSC Willed Body Program. Inclusion criteria include healthy donor eyes that are free from any ocular diseases and other neurodegenerative conditions. A total of six to eight explants or four equal retinal quadrants were isolated from each healthy retina and placed on Transwell Permeable 6.5 mm inserts or in 12 well plates with the RGC layer facing up. The retinal explants and quadrants were cultured for 7 days under neurotrophic factors deprivation conditions and treated either with DMSO or Torin 2 (100 μM) in an explant medium consisting of phenol red-free Neurobasal A with 2% B-27, 1% N-2, 0.8 mM L-glutamine, 100 U/mL penicillin, and 100 µg/mL streptomycin (all reagents were purchased from Thermo Fisher Scientific, Waltham, MA, USA). Following the 7-day treatment, the retinal explants were fixed with 4% PFA for 24 hr at 4°C and then washed 3 times in 1X PBS. All explants were blocked with blocking buffer (10% goat serum, 0.2% Triton X-100 in 1X PBS) for at least 2 hr at room temperature and then incubated with RBPMS antibody for three days at 4°C. Following washing 3 times in 1X PBS, the retinal explants were further incubated with a corresponding secondary antibody (goat anti-rabbit 568, 1:500; Invitrogen) for 2 hr at room temperature. The retinal explants were rinsed 3 times in 1X PBS and mounted with DAPI mounting solution on glass slides. Images were captured using a Keyence microscope, and RBPMS-positive cells were counted using ImageJ software.

### Treatment of mice with Torin 2

Three-month-old Mt-Keima mice were intravitreally injected with Torin 2 (1 mM prepared in DMSO) in one eye, and the contralateral eye was given DMSO as a control under isoflurane anesthesia. Mt-Keima mice were euthanized 24 hrs after the treatment, and mitophagy flux was measured in ON *in vivo*. In C57BL/6J mice, IOP was elevated for 7 weeks using weekly periocular injections of Dex and then intravitreally injected in one eye with Torin 2 (1 mM prepared in DMSO) and the contralateral eye with DMSO. Three weeks after Torin 2 treatment under sustained IOP elevation conditions, mice were euthanized, and the structural and functional integrity of both the RGCs and the axons was assessed.

### Adeno-Associated Viral 2 (AAV2) vector injections

AAV2 vectors expressing empty/null or Cre or Parkin under the control of a synapsin (Syn) promoter were purchased from VectorBuilder. For RGC-specific conditional knockdown of *Atg*5, three-month-old ATG-5flox/flox mice were injected intravitreally one eye with AAV2-Syn-Cre, and the contralateral eyes were injected with AAV2-Syn-empty vector using a Hamilton syringe fitted with a sterile 33-gauge needle. Following 6 weeks post-injection, mice were euthanized, and RGC-specific *Atg*5 knockdown and glaucoma phenotypes were examined by immunostaining on retinal cross-section. For RGC-specific Parkin overexpression, three-month-old C57BL/6J mice were injected intravitreally one eye with AAV2- Syn-Parkin and the contralateral eye with AAV2-Syn-empty using a Hamilton syringe fitted with a sterile 33-gauge needle. Following 2 weeks post-injection, IOP was elevated bilaterally in those mice for 10 weeks using Dex periocular injections. At the end of the study period, mice were euthanized, and glaucoma phenotypes were examined.

### Antibodies and reagents

Antibodies and reagents were purchased from the following sources: RBPMS (catalog # 118619, Gene Tex), TOMM20 (catalog # sc-17764 from Santa Cruz and catalog # 11802 from Proteintech Inc), PINK1 (catalog # BC100-494, Novus), Parkin (catalog # sc-32282, Santa Cruz), CoxIV (catalog # 11242-1, Proteintech), SOD2 (catalog # ab16956, Abcam), 8-OHdG (catalog # sc-66036, Santa Cruz), p62 (catalog # PM045, MBL), LC3 (catalog # GTX127375, Gene Tex). Goat anti-Rabbit IgG (H+L) Cross-Adsorbed Secondary Antibody, Alexa Fluor™ 568 (catalog # A-11011, Invitrogen), Goat anti-Rabbit IgG (H+L) Highly Cross-Adsorbed Secondary Antibody, Alexa Fluor™ 488 (catalog # A-11034, Invitrogen), Goat anti-Rabbit IgG (H+L) Cross-Adsorbed Secondary Antibody, Alexa Fluor™ 680 (catalog #A-21076, Invitrogen), Goat anti-Mouse IgG (H+L) Highly Cross-Adsorbed Secondary Antibody, Alexa Fluor™ 568 (catalog # A-11031, Invitrogen), Goat anti-Mouse IgG (H+L) Cross-Adsorbed Secondary Antibody, Alexa Fluor™ 488 (catalog # A-11001, Invitrogen). Dexamethasone, Micronized, USP (Spectrum, DE121), Torin 2 (Millipore Sigma, SML 1224), 16% PFA Aqueous Solution EM Grade (Electron Microscopy Sciences, 15710-S), Aqueous Glutaraldehyde EM Grade, 10% (Electron Microscopy Sciences, SKU 16120), Sodium Cacodylate Buffer, Electron Microscopy Sciences (catalog # 102090-962, Avantor), DAPI (VECTASHIELD Antifade Mounting Medium, Vector Laboratories).

### Statistical Analysis

GraphPad Prism v10.0 software was used for statistical analysis. N refers to the total number of eyes used for the study. Data was represented as mean±SD. For the comparison of two different study groups, we used a 2-tailed Student’s t-test, and for the comparison of three study groups, 2-way ANOVA with multiple comparisons was used. A value of *p < 0.05* was considered statistically significant.

### Scientific illustration

BioRender was used to create the illustrations for this article.

### Study approval

Animal studies were conducted in compliance with the Association for Research in Vision and Ophthalmology Statement on the Use of Animals in Ophthalmic and Vision Research and the Institutional Animal Care and Use Committee regulations of the University of North Texas Health Science Center (approved protocol: IACUC2018-0032 and IACUC2020-0026) and University of California Irvine (approved protocol: AUP-22-159).

## Data Availability Statement

All the datasets used and/or analyzed in the present study is available from the corresponding author on reasonable request.

## Authors Contributions

P.M and G.S.Z. designed research studies, analyzed data, performed experiments and wrote the manuscript. BRK has conducted experiments involving Tg. Cre-MYOC^Y437H^ mouse model of POAG. BK performed human retinal explant experiments as well as PPD imaging. KK assisted with TEM imaging and immunostaining. JCM, SY, and RBK participated in Torin 2-related experiments on a GC-induced mouse model of glaucoma. AFC assisted in key experiments and provided the human donor tissues as well as reagents. All authors discussed the results and implications and commented on the manuscript at all stages. All authors read and approved the final manuscript.

## Acknowledgments

These studies were supported by the National Institutes of Health; EY026177 (GZ), EY028616 (GZ) and K99EY032982 (PM) and the Bright Focus Glaucoma Foundation; G2022004F (PM) and the unrestricted startup funds from the University of Missouri, Columbia (PM). The authors also acknowledge the support from the NIH grant EY034238 and from an unrestricted grant from Research to Prevent Blindness to the Gavin Herbert Eye Institute at the University of California, Irvine.

The authors would like to thank Charles Kiehlbauch (UNTHSC), Dr. Dorota Stankowska (UNTHSC), Dr. Denise Inman (UNTHSC), Dr. Raghu Krishnamoorthy (UNTHSC), Sherri Feris (UNTHSC), Stacy Curry (UNTHSC), Linya Li (UCI) and Rowan Fink (MU), Anne-Marie Brun (UNTHSC) and Chantal Allamargot (University of Iowa) for assistance with some experiments.

