## supplemental for "Genetic and pharmacological correction of impaired mitophagy in retinal ganglion cells rescues glaucomatous neurodegeneration"

**Supplementary figure 1:**

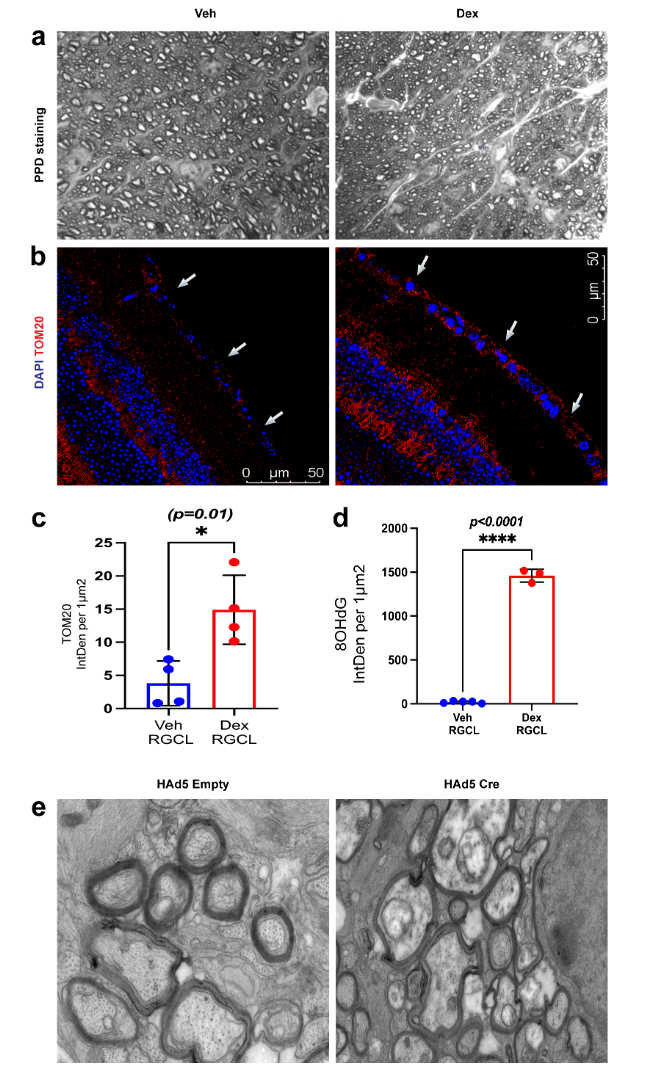

**OHT leads to dysfunctional mitochondrial accumulation: a)** PPD images of ON cross sections showed severe axonal degeneration, **b and c)** an increased expression of TOM20 and its quantitative analysis, and **d)** quantitative analysis of 8-OHdG in RGCL of Dex-induced OHT eyes compared to Veh-injected eyes. **e)** TEM imaging on ON cross sections from Tg.Cre-MYOC^Y437H^ mice injected with HAd5-Cre and HAd5-empty vectors. (n=4 in each group, two-tailed unpaired t-test).

**Supplementary figure 2:**

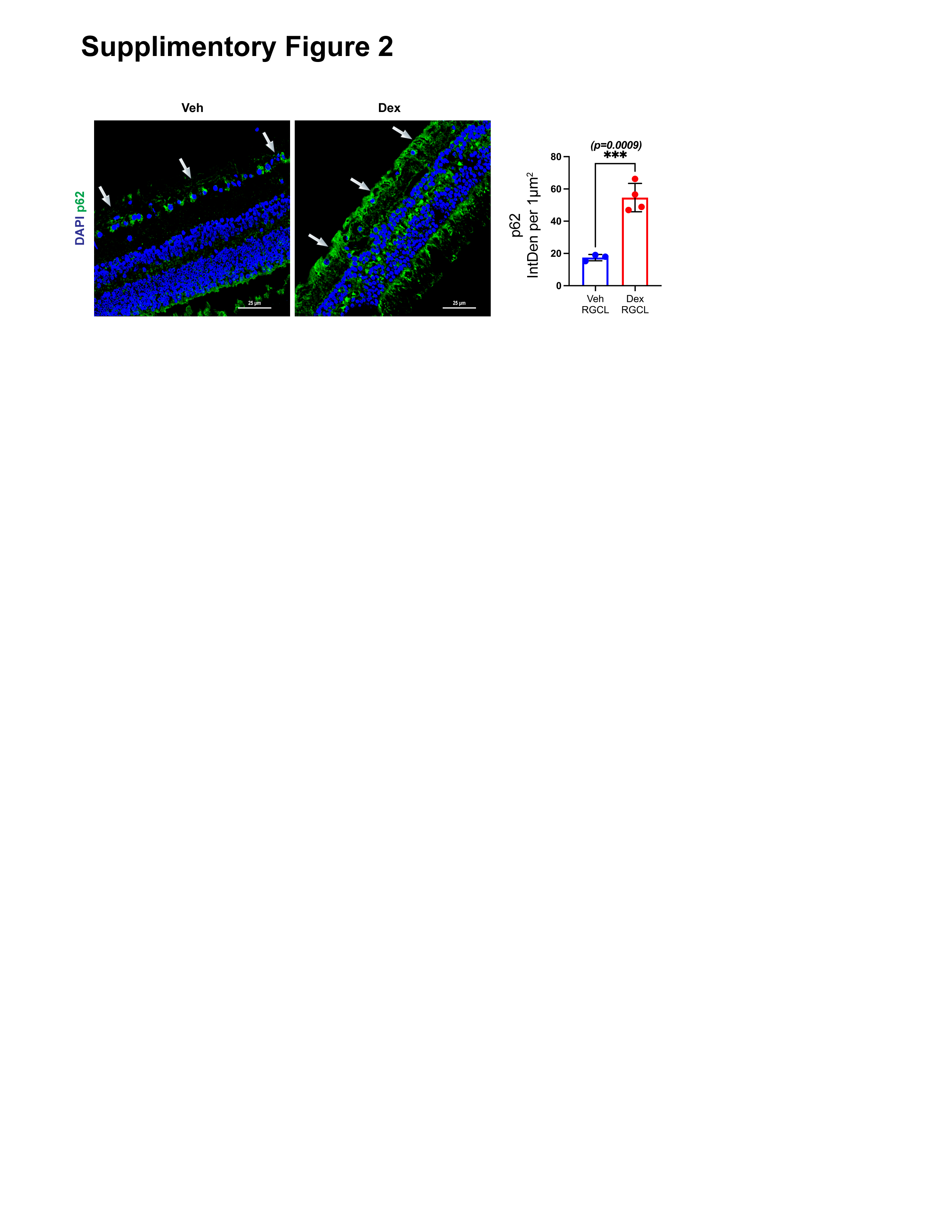

**OHT leads to impaired mitophagy:** Based on immunostaining and quantitative analysis data, p62 expression was increased in the RGCL of GC-induced mouse models of glaucoma (n = 4 in each group, two-tailed unpaired t-test, and arrows represent RGCL).

**Supplementary figure 3:**

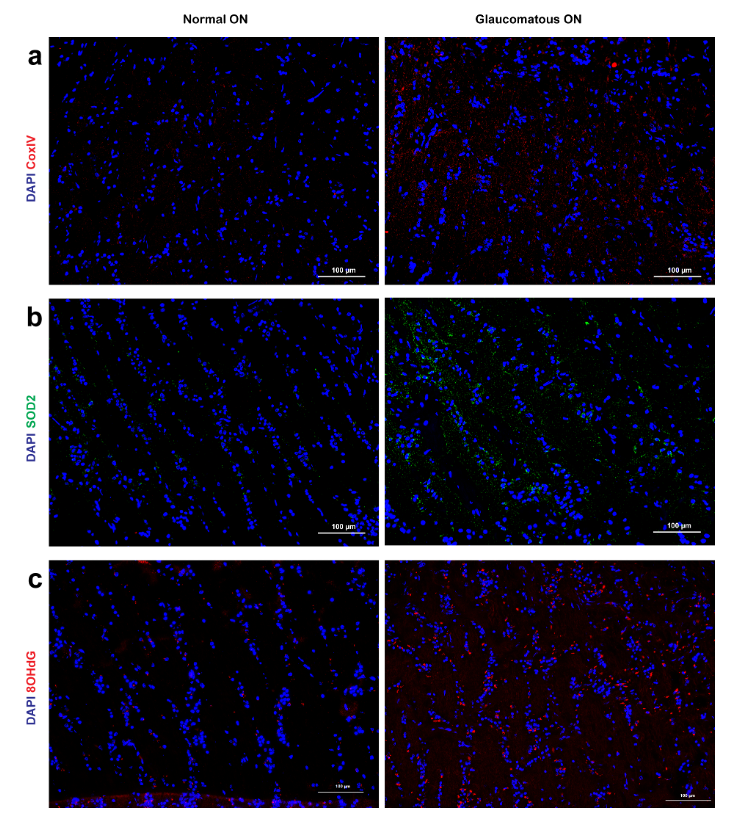

**Human glaucoma pathology is associated with dysfunctional mitochondrial accumulation and oxidative stress:** An increased expression of **a)** mitochondrial marker CoxIV, **b)** mitochondrial oxidative stress marker SOD2, and **c)** oxidative DNA damage marker 8-OHdG in age matched human glaucomatous ON compared to the normal ON.

**Supplementary Table 1. List of human donor tissue and ocular pathology**

| **Donor Tissue ID** | **Ocular Pathology** | **Age** | **Race** | **Sex** | **Cause of Death** |
| --- | --- | --- | --- | --- | --- |
| 842-09 | Normal; IOL sx 5 years ago | 82 | C | F | Stroke-CVA |
| 528-10 | Normal | 71 | C | M | Sudden cardiac event |
| 301-11 | Normal | 85 | C | M | Pancreatitis |
| 839-10 | Normal | 84 | C | F | Cerebral vascular accident |
| 371-11 | Normal | 85 | C | M | Probably heart disease |
| 241-10 | Normal | 77 | C | M | Cardiac event, multiple myeloma |
| 694-10 | Normal | 76 | C | M | Congestive heart failure |
| 236-11 | Normal | 74 | C | M | Respiratory arrest |
| 49-11 | Glaucoma | 78 | C | M | Acute cardiac crisis |
| 140-11 | Glaucoma | 76 | C | F | Multi system organ failure |
| 1109-10 | Glaucoma | 80 | C | M | Pneumonia |
| 115-11 | Glaucoma open angle | 81 | C | M | End stage Alzheimer’s |
| 608-10 | Glaucoma | 76 | C | M | Lung CA |
| 936-10 | Glaucoma | 82 | C | M | COPD |
| 616-10 | Glaucoma, | 83 | C | F | COPD |
| 284-10 | Glaucoma | 67 | C | M | Liver CA |

C, Caucasian; F, Female; M, Male; COPD, Chronic obstructive pulmonary disease; CA, Cancer; CVA, Cerebrovascular Accident.
